# Formulation of FMD vaccine in Naloxone/Alum mixture: A potency study

**DOI:** 10.1101/2022.07.19.500605

**Authors:** Danasadat Alemalhoda, Farzam Ajamian, Akbar Khorasani, Setareh Haghighat, Mona Mahdavi Amreie, Fatemeh Sadat Sarkeshikzadeh Moghadas, Fatemeh Asgarhalvaei, Alireza Kalantari, Mehdi Mahdavi

**Author notes:** **Correspondence:** Alireza Kalantari, department of genetics, university campus 2, university of guilan, guilan, iran.

## Abstract

**Background:** Current Foot-and-Mouth disease (FMD) vaccines are commonly formulated in the alum adjuvant. Several studies showed that this form of vaccine, although able to control the infection, lacks the ability to eradicate the infection completely.

**Objectives:** In this study, the FMD vaccine was formulated in a naloxone/alum mixture as an adjuvant, and its potency was compared with the alum-formulated vaccine.

**Methods:** Experimental Balb/c mice were immunized with a commercial vaccine or naloxone/alum-based vaccine two times, subcutaneously at two-week intervals. Lymphocyte proliferation was assessed using BrdU, and IL-4, IFN-γ, and TNF-α cytokines, specific total IgG as well as IgG1/IgG2a were evaluated by ELISA. In addition, the gene expression profiles of IL-12, IL-17, and IFN-γ cytokines were determined by real-time PCR.

**Results:** Results showed that mice immunized with the FMD vaccine formulated with the naloxone/alum mixture exhibited a tiny increase in the production of IFN-γ and TNF-α cytokines compared to the routine vaccine. However, mice immunized with the FMD vaccine formulated with the naloxone/alum mixture revealed a significant increase in the expression of IL-12, IL-17, and IFN-γ cytokines compared to those immunized with the routine vaccine. In addition, the novel formulation led to increased production of specific total IgG in comparison with the routine vaccine.

**Conclusions:** Our findings suggest that naloxone formulated in the FMD vaccine could improve cellular and humoral immune responses. However, the effect of this formulation on the efficacy of vaccine is remained to be clarified in future studies.

## 1. Introduction

Foot-and-Mouth disease (FMD) is a highly contagious and harmful disease affecting cloven-hoofed animals, which causes great economic losses worldwide (1). FMD is endemic in many regions of the world, such as parts of Asia, Africa, South America, and the periphery of the European Union (2). Clinical effects of FMD in cattle cause loss of anorexia, fever, and severe weight loss (3). There are seven serotypes of FMD virus (FMDV), including O, A, C, SAT 1, SAT 2, SAT 3, and Asia 1, in which type O is the most prevalent serotype. Serotypes A, O, and Asia1 are present in Iran (4-6). Vaccination with high-quality vaccines is one of the most important strategies for controlling and eradicating FMD, especially in enzootic areas. The advent of mass vaccination significantly decreased the morbidity or mortality of newborns and adults (7).

Adjuvants are essential components to induce an appropriate response in inactivated vaccines. A suitable adjuvant can potentially control infections and eliminate virus-infected cells. Alum, as the most used adjuvant, stimulates T helper 2 (Th2) responses but cannot induce the Th1 responses required to control intracellular pathogens. The Th2 immune response alone lacks the ability to protect against diseases fully; on the other hand, Th1 immunity is essential for protection against infectious diseases, such as FMDV (7-9).

Traditional FMD vaccines are often formulated in the alum adjuvant. This form of vaccine, although having the ability to control the infection, cannot completely eradicate the infection. Importantly, a variety of studies showed that this adjuvant is not a completely ideal adjuvant for confronting infections such as FMD infection, because of the fact that carrier livestock can transmit the virus to other livestock (10, 11). Therefore, there is a need to replace or optimize the alum adjuvant to induce cellular immune responses. A wide variety of studies have shown that naloxone (NLX), an opioid receptor antagonist, has adjuvant effects (12, 13) and can shift the immune response toward the Th1 pattern (1, 2).

Cell-mediated immunity plays a critical role in host resistance to intracellular infections induced by bacteria, parasites, or viruses (2). In addition, NLX can activate p38 mitogen-activated protein kinases (MAPK), inhibit nicotinamide adenine dinucleotide phosphate (NADPH) oxidase and cause an increase in pro-inflammatory cytokines that results in inflammation (14, 15). Cytokines have an important role in controlling viral and bacterial infections, which induce their effects by modulating immune responses and directing immune response patterns in exposure to pathogens and finally defeat pathogens (16). The level of pro-inflammatory cytokines increases in the initial stages of the disease, which helps polarization toward Th1 pattern (16, 17). In recent years, there have been many improvements in developing of vaccine candidates for FMD. The limitation of current FMD vaccines is the lack of an appropriate adjuvant to stimulate both cellular and humoral immunity.

According to experiences with NLX as an adjuvant in the polarization toward the Th1 pattern (1, 2, 18). Herein we hypothesized that FMD formulated with the alum/NLX mixture, as an adjuvant, may modify immune response patterns and provide more potent cellular and humoral immune responses.

## 2. Materials and methods

### 2.1. Production and purification of FMD virus (FMDV)

The FMDV serotype O2010/IR was propagated in BHK21 cell culture, harvested, centrifuged to remove cell debris, and then concentrated with 8% polyethylene glycol 6000. The virus concentration titer was measured by the TCID50 method and inactivated by 4 mM w/v Binary Ethyleneimine (BEI) for 30 hours at 30°C. Lastly, 2 mM sodium thiosulfate was added to neutralize and remove residues of BEI.

### 2.2. Vaccine formulation

For this purpose, inactivated FMDV particles were formulated in alum and alum/NLX mixtures with the standard protocol. Each vaccine formulation used 10 µg inactivated FMD virus type O2010/IR as an antigen payload per dose. Each dose equal 100 µl of the vaccine was used for immunization purposes. For the alum/NLX mixture, 120 µg of NLX (6 mg/kg) were used in a total volume of 100 µl at the injection time.

### 2.3. Mice and immunization protocol

Six-to eight-week-old inbred BALB/c male mice (n=35) were obtained from Razi Vaccine and Serum Institute (Table 1). The mice were divided into five groups. Experimental mice were immunized subcutaneously two times with 10 µg of the FMD vaccine formulated in the alum or alum/NLX mixture at two-week intervals with matched control groups. All experiments were in accordance with the Animal Care and use Protocol of Pasteur Institute of Iran.

**Table 1.**
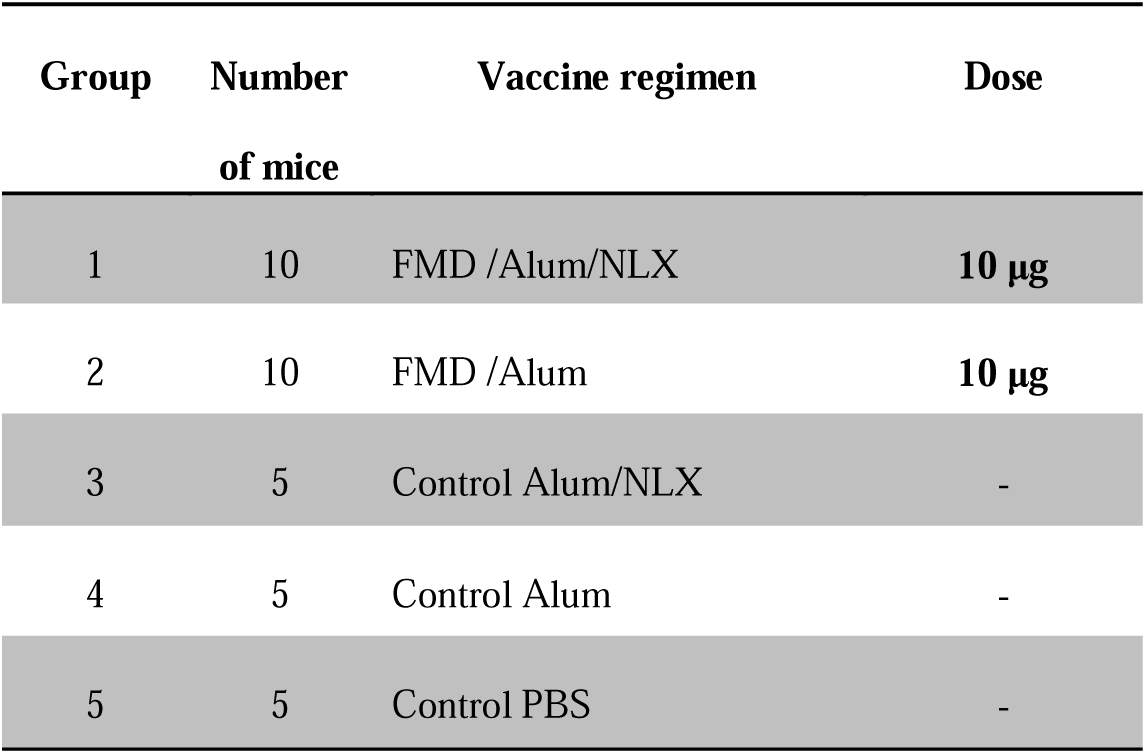
Experimental groups and immunization

| Group | Number<br>of mice | Vaccine regimen | Dose |
| --- | --- | --- | --- |
| 1 | 10 | FMD /Alum/NLX | 10 µg |
| 2 | 10 | FMD /Alum | 10 µg |
| 3 | 5 | Control Alum/NLX | - |
| 4 | 5 | Control Alum | - |
| 5 | 5 | Control PBS | - |

### 2.4. Serum sampling

Ten days after the last immunization, blood samples were collected by retro-orbital puncturing of experimental mice. Sera were separated by centrifugation and stored at −20 °C for antibody assay.

### 2.5. Lymphocyte proliferation by BrdU method

Ten days after the last vaccination, the mice were dislocated, and the spleens were placed in sterile conditions and suspended in spleen washing buffer containing PBS with 10% FBS, Pen/strep and 4 mM L-Glutamine. Cell suspension was centrifuged at 2500 rpm for 5 minutes at 4 □C. Then, the supernatant was discarded, and 5ml mouse RBCs lysis buffer was added to the cells, 5 ml of spleen washing buffer was added to each tube after 5 minutes. Centrifugation was conducted as explained in the previous step. The cell suspension was prepared and adjusted to 4×10^6^ cells/ml in RPMI-1640 with 10% FBS, 4 mM L-glutamine, 25 mM 4-2-Hydroxyethlypiperazine-1-2-Ethanesulfonic Acid (HEPES), 0.1 mM non-essential amino acid, 1 mM sodium pyruvate, 50 µm 2ME, 100 µg/ml streptomycin and 100 IU/ml penicillin (Gibco, Germany). Then, 100 µl of samples with 4×10^5^ cells were cultured in 96 well culture plates (in triplicate) and stimulated with 5 µg/ml of the FMD antigen as a recall antigen. In addition, Con-A (1μg/ml; Sigma, USA) and un-stimulated wells were used as positive and negative controls, respectively. The plates were incubated at 37°C in a 5% CO_2_ humid incubator for **7**2 hours. Then, 20 μl of 10X 5-Bromo-2’-deoxyuridine (BrdU) (Roche, Germany) per well was added, and culture was continued for 24 hours. Thereafter, the plates were centrifuged at 300×g for 10 minutes, and the culture medium was removed and dried at 60 °C for 30 minutes. In the next step, 200 μl of fixation/denaturation buffer was added to the wells and incubated at room temperature (RT) for 30 minutes. Subsequently, the cells were permeabilized, and then 100 µl of anti-BrdU antibody was added to each well and incubated for 90 minutes at RT. The plates were washed six times with PBS, and 100 µl of Tetra-Methyl Benzidine (TMB, Roche, Germany) solution was added to each well, and the reaction was continued for 10 minutes in the dark. Finally, 100 µl of 2N H_2_SO_4_ was added to stop the reaction, and absorbance eas measured in 450 nm wavelength by an ELISA reader.

### 2.6. IL-4, IFN-γ and TNF-α cytokine assay

Spleen cell suspension containing 4×10^6^ cells/ml was cultured in 24-well plates and stimulated with 5 µg/ml of the FMD antigen. The plates were then incubated in a CO2 incubator at 37°C for 72 hours. After the incubation period, the spleen cell suspensions were collected. The suspensions were used to assess the cytokine levels, including IL-4, IFN-γ, and TNF-α, by using Mabtech ELISA kits (Sweden) according to the manufacturer’s instruction. The pg/ml ratio of each sample was calculated according to the standard curve, and the absolute cytokine production of each mouse was used for statistical analysis.

### 2.7. Real-time PCR of IL-12, IL-17, and IFN-γ cytokine gene expression

#### 2.7.1. IL-12, IL-17 and IFN-γ cytokine primers

The sequences of IFN-γ, IL-4, and IL-17 cytokines and accession numbers were selected after searching the publicly available databases (Gene Bank) (accession numbers NC_000076.6, NC_000077.6, and NC_000067.6, respectively). Primers were designed using Primer Express 3.0 software. Table 2 indicates the details of all oligonucleotide primer sequences, primer lengths, and accession numbers.

**Table 2.**
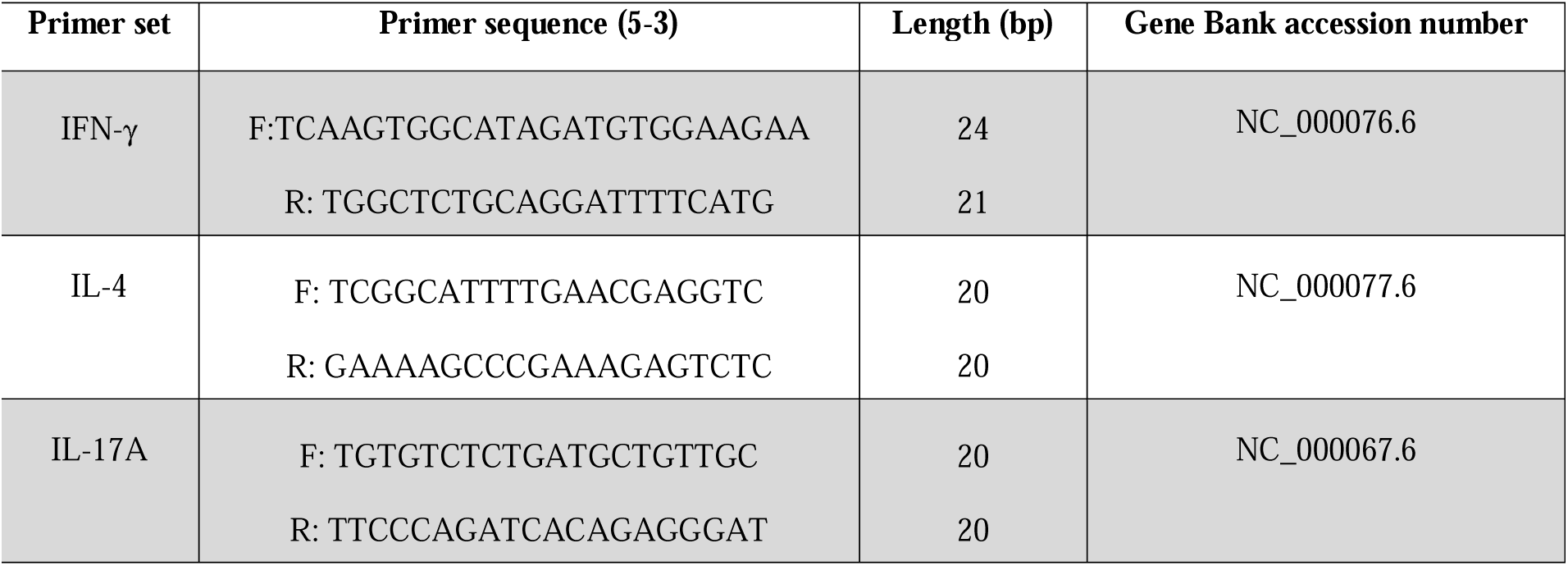
Sequences of oligonucleotide primers used for quantitative real-time PCR

#### 2.7.2. RNA extraction and cDNA synthesis

Spleen cells of experimental mice were cultured for 12 hours with or without the antigen (the optimized time calculated according to our prior experiences), and then total cellular RNA was extracted from cells by QIAzol Lysis Reagent (QIAGEN, USA) according to the manufacturer’s instruction. cDNA was synthesized using random hexamers, and then cDNA was stored at −20 C until use.

#### 2.7.3. Real-time PCR

Real-time was performed with RealQ-PCR 2x Master Mix Kit (GENE ALL company) according to the manufacturer’s instructions. The reaction used for the One-Step rRT-PCR was as follows: 5 µl SYBR® Premix EX Taq II (2X), 1.0 µl of each forward and reverse primer, 1.0 µl of the template, and 2.0 µl dH2O. The amplification was carried out at the following temperature cycle: The cycling parameters were 95 °C for 15 min, followed by 40 cycles consisting of 95 °C for 15 seconds, 62°C for 20 seconds. Each sample was tested in duplicate. PCR amplification was carried out in the Thermo cycler Rotor GeneQ (Qiagen, Germany). Differences in gene expression were calculated by the comparative ΔΔCT method (19).

##### 2.7.3.1. Specific total IgG and IgG1/IgG2a isotypes

Ten days after the last immunization, the levels of total antibodies and IgG1, IgG2a isotypes were assessed by indirect ELISA. For this purpose, 100µl of inactivated FMDV (10µg/ml) in PBS were coated in 96-well ELISA Maxisorp plates (Greiner, Germany), followed by overnight incubation at 37°C. In the next step, the wells were washed three times with PBS containing 0.05% Tween 20 (washing buffer) and blocked with blocking buffer containing 3% skimmed milk in PBS buffer for one hour at 37°C. The washing step was carried out according to the previous step. Serial dilutions of sera (1/10 to 1/1280) were prepared in dilution buffer containing 1% BSA in PBS with 0.05% Tween 20, and 100µl of each dilution was added to each well and incubated at 37°C for 2 hours. After incubation, the wells were washed five times with washing buffer, and 100µl of 1/10000 dilution of anti-mouse HRP conjugate (Sigma, USA) in dilution buffer was added and incubated for 2 hours at 37°C. After that, the wells were washed, as explained in the previous step. Then, 100µl of TMB substrate was added to all wells and incubated for 30 minutes in the dark. The reaction was stopped by adding 100 μl of 2N H_2_SO_4,_ and color density was measured at 450nm wavelength with an ELISA plate reader. In order to evaluate the levels of specific IgG1 and IgG2a, the goat anti-mouse IgG1, IgG2a secondary antibodies (Sigma, USA) were used in the step following sampling and incubated at 37°C for 2 hours. Other steps were conducted as described in total antibody ELISA.

### 2.8. Statistical analysis

The data are expressed as Mean±SD of triplicate experiments, and statistical analysis was carried out using Graph pad prism software (V 6.01) using Mann Whitney U method. P-value less than 0.05 was considered significant between experimental groups.

## 3. Results

### 3.1. Lymphocyte proliferation assay

The spleen cell proliferation responses were demonstrated among vaccinated and control groups, as illustrated in Fig. 1. The results showed that injection of the vaccine formulated in the alum/NLX mixture caused a significant increase in proliferation responses as compared with alum/NLX and PBS control groups (P<0.0107). Immunization with FMD-alum/NLX showed an approximately 11% increase in lymphocyte proliferation as compared with the FMD-alum group but statistically was not significant (P=0.6839).

**Figure 1.**
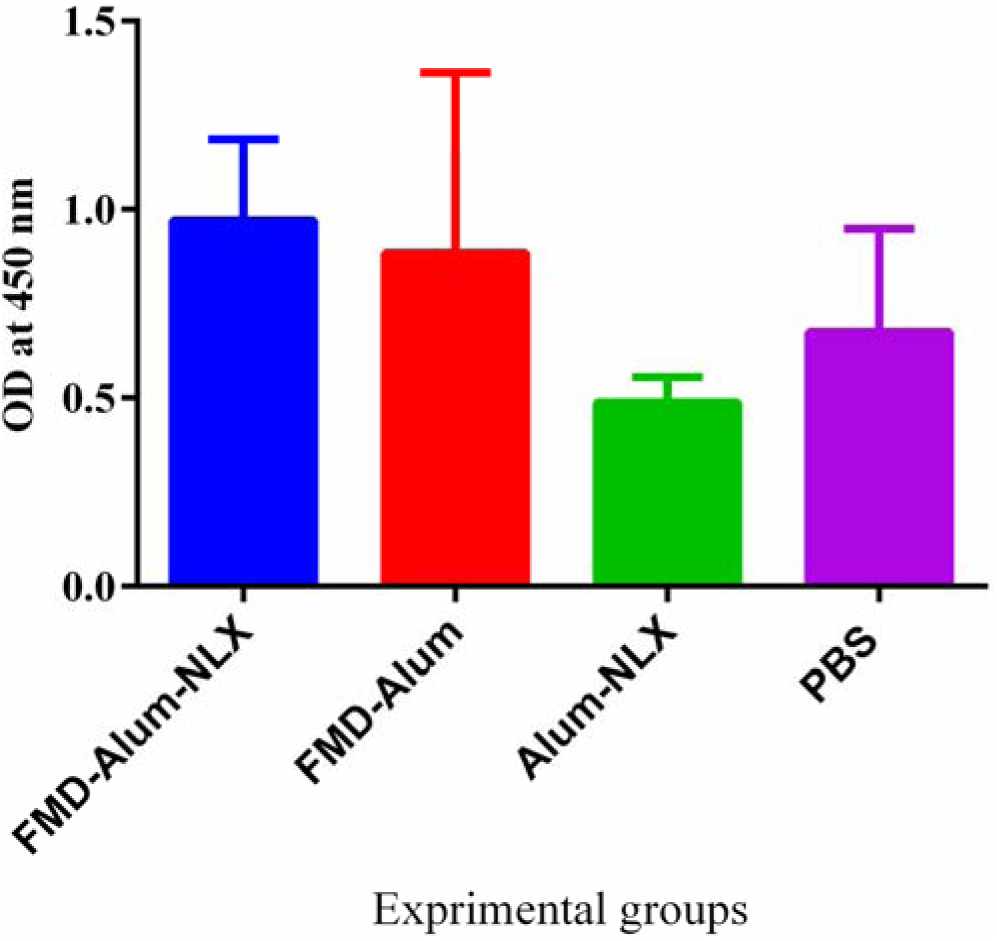
lymphocyte proliferation in the experimental groups. The results show that injection of the vaccine formulated in alum/naloxone mixture caused a significant increase in proliferation responses compared to the alum/NLX and PBS control groups (P<0.0107). And also a tiny increased versus routine vaccine. Data are shown as mean ± SD.

### 3.2. IL-4, IFN-γ, and TNF-α cytokines ELISA

Results from the IL-4 cytokine assay showed that immunization with FMD-alum/NLX and FMD-alum vaccines demonstrated a significant increase in the IL-4 levels as compared with the control groups (P<0.0091) (Fig. 2a). No significant difference was observed between FMD-alum/NLX and FMD-alum vaccine groups in the secretion of the IL-4 cytokine (P=0.2870).

**Figure 2.**
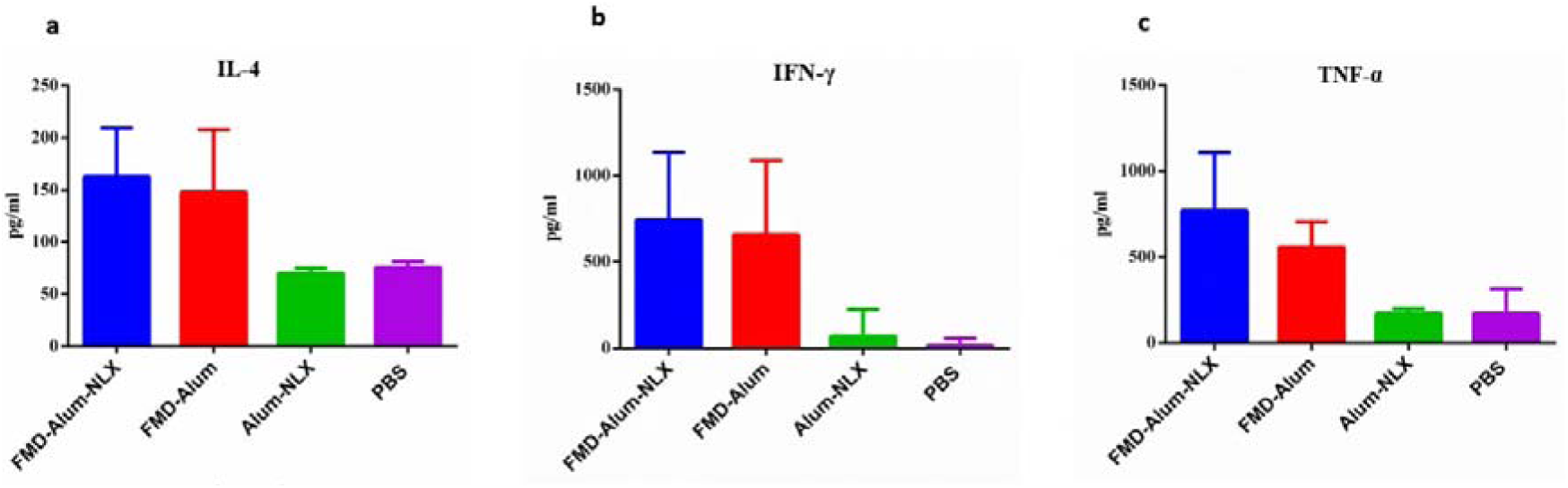
IL-4, IFN-γ, and TNF-α cytokine levels in the experimental groups. **a)** The result of IL-4 cytokine in the experimental groups shows that immunization with FMD-alum-naloxone and FMD-alum vaccines demonstrated a significant increase in IL-4 production as compared with control groups (P<0.0091). **b)** IFN-γ levels exhibit a significant increase in the FMD-alum-naloxone and FMD-alum experimental groups compared to the control groups (P<0.0378). Novel formulation shows a slight increase in the IFN-γ levels compared to the routine vaccine. Data are shown as mean ± SD. **c)** The result from TNF-α cytokine indicates that immunization with FMD-alum-naloxone and FMD-alum significantly increases TNF-α cytokine compared with the control groups (P<0.0167). In addition, immunization with a novel vaccine shows an about 38% increase in the TNF-α cytokine secretion compared to the FMD-alum group.

Mice immunized with the FMD-alum/NLX, and FMD-alum groups showed a significant increase in the IFN-γ levels compared to the control groups (P<0.0378). In addition, mice immunized with FMD-alum/NLX showed a 13% increase in the IFN-γ cytokine secretion compared with the FMD-alum group but were not significantly different (P= 0.3799) (Fig. 2b).

Mice immunized with FMD-alum/NLX, and FMD-alum showed a significant increase in the TNF-α cytokine production as compared with the control groups (P<0.0167). In addition, mice immunized with FMD-alum/NLX showed about 38% increase in the TNF-α cytokine secretion compared with the FMD-alum group (P= 0.1216) (Fig. 2c).

### 3.3. Real-time PCR of IL-12, IL-17, and IFN-γ

Real-time PCR analysis of IFN-γ (Fig. 3a), IL-12 (Fig. 3b), and IL-17 (Fig. 3c) gene expression displayed that mice immunized with the FMD-alum/NLX vaccine had a significant increase in these cytokines as compared with the FMD-alum as well as control groups (P<0.005).

**Figure 3.**
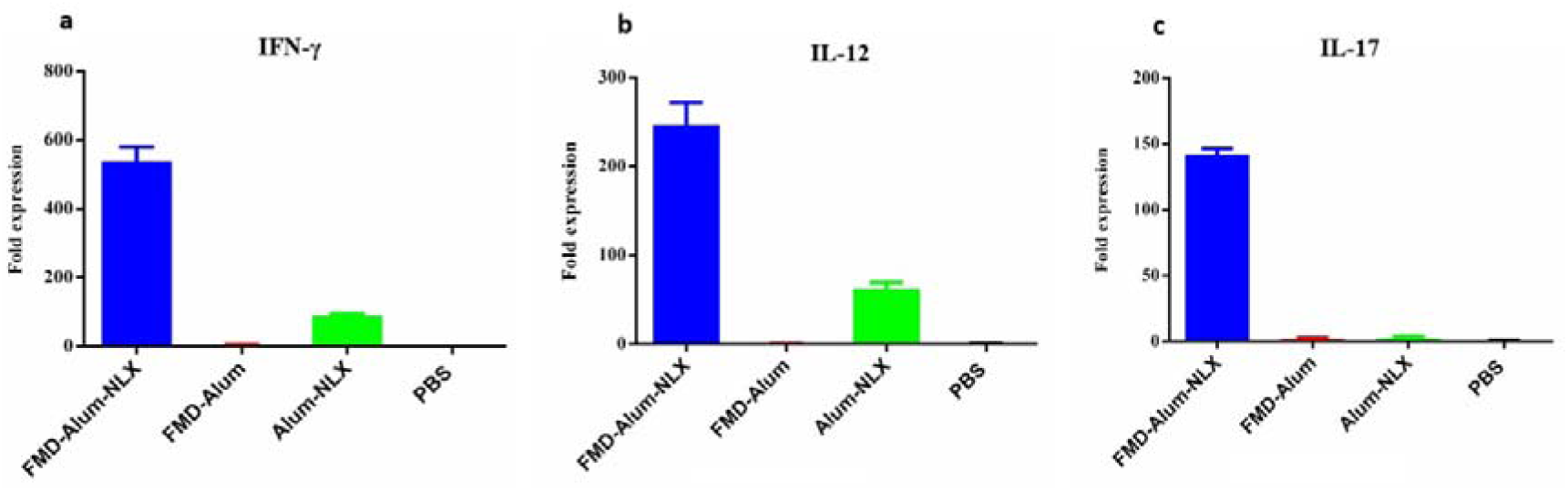
the mRNA expression level of the IFN-γ, IL-12, and IL-17 cytokine in the experimental groups. **a)** IFN-γ gene expression shows a significant increase in the IFN-γ expression in mice immunized with the FMD-naloxone-alum vaccine compared to the FMD-alum and control groups (P<0.005). Data are shown as mean ± SD. **b)** The results show a significant increase in the IL-12 cytokine in the mice immunized with the FMD-naloxone-alum vaccine compared to those immunized with the FMD-alum and control groups (P<0.005). Data are shown as mean ± SD. **c)** The results show that mice immunized with the FMD-naloxone-alum vaccine have a significant increase in the IL-17 cytokine as compared with the FMD-alum and control groups (P<0.005).

### 3.4. Total IgG and specific IgG1/IgG2a isotypes

Our findings showed that all immunized groups (FMD-alum/NLX and FND-alum) significantly increased total antibodies in comparison with the control groups (dilutions of 1/10 up to 1/320, P<0.0359) (Fig. 4a). In addition, mice immunized with FMD-alum/NLX vaccine increased specific total IgG as compared with the FMD-alum group at dilutions of 1/10 and 1/20 (P=0.0787 and P=0.0543, respectively).

**Figure 4.**
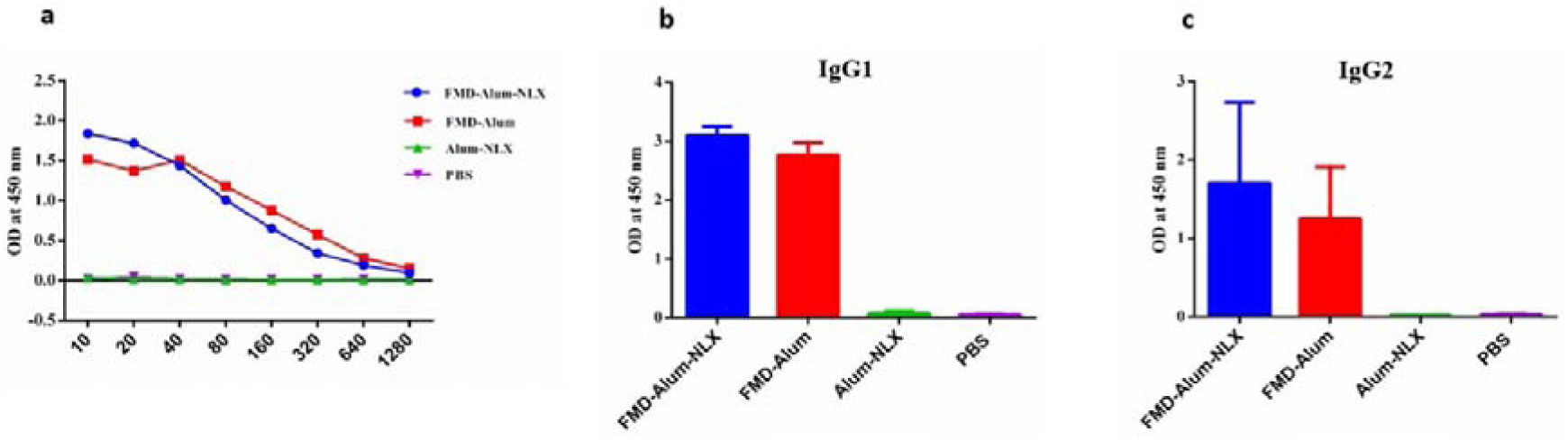
Serum-specific total IgG, IgG1, and IgG2a antibody levels in the experimental groups 10 days after the last shot. **a)** Our findings show that immunization with the FMD-naloxone-alum vaccine increased specific total IgG compared to the FMD-alum group at dilutions of 1/10 and 1/20 (P=0.0787 and P=0.0543, respectively). **b)** There is a significant increase in the IgG1 response of the vaccine adjuvanted with the naloxone-alum mixture compared to the FMD-alum group (P=0.0019). Data are shown as mean ± SD. **c)** Mice administered with FMD-naloxone-alum showed an increase in the production of IgG2a as compared with the FMD-alum group (P=0.0622). Data are shown as mean ± SD.

Results from specific IgG1 (Fig. 4b) and IgG2a (Fig. 4c) showed that mice immunized with FMD-NLX-alum and FMD-alum vaccines significantly increased specific IgG1 and IgG2a isotypes as compared with the control groups (P<0.0001). There was a significant increase in the IgG1 response in the mice immunized with the vaccine adjuvanted with alum/NLX mixture compared with the FMD-Alum group (P=0.0019). Additionally, mice immunized with FMD-alum/NLX increased the IgG2a response compared to the FMD-alum group (P=0.0622).

## 4. Discussion

Adjuvants are a crucial component of vaccines that improve vaccine efficacy by increasing the speed and magnitude of immune response development and decreasing the amount of antigens (20). The selection of adjuvants is an important step in vaccine formulation for induction of humoral and cellular immunity as well as protection. In addition, the selection of an appropriate adjuvant is one of the most important factors in determining the efficacy of vaccines (21). Alum is a routine adjuvant used in commercial and FMD vaccines (21). Alum, although inducing Th2-type responses in animals and humans (22, 23), lacks the ability to stimulate Th1 responses. Appropriate immune responses against viral pathogens, such as FMD, are induction of both Th1 and Th2 immune responses (24). Importantly, Th1 responses, especially the IFN-γ response, are essential for completely cleaning infected cells, and such responses prevent carrier formation in the cattle’s for FMD infection (24, 25).

Nowadays, adjuvants should be optimized for increased efficiency and decreased toxicity of vaccines. The alum-based FMD vaccine should be optimized for better performance due to its limitation in controlling FMD infection (23). NLX, an adjuvant that has recently been studied, was demonstrated to have great potential in the induction of the Th1 pattern and synergistic effects with alum in the induction of cellular and humoral immune responses (18, 24, 26). Here, the adjuvant activity of the alum/NLX mixture was evaluated against the FMD vaccine model, and we hypothesized that the formulation of the FMD vaccine in the alum/NLX mixture might improve its potency for induction of higher levels of cellular and humoral immune responses.

Lymphocyte proliferation assay shows that the FMD vaccine formulated in the mixture of alum/NLX showed a tiny increase in lymphocyte proliferation compared with the alum-based vaccine. Previous studies on NLX/alum mixture in various vaccine models showed a synergistic effect on lymphocyte proliferation (27, 28) which we achieved very weak in the FMD vaccine model. This may show that the adjuvant effect of alum/NLX mixture may vary according to the antigen nature, and this property may not be general for all immunogens.

We also assessed the secretion of IL-4, IFN-γ, and TNF-α cytokines. Results from the present study indicated that the mice immunized with the FMD-alum/NLX vaccine showed a tiny increase in the levels of IL-4 and IFN-γ but a higher level of TNF-α as compared with those immunized with the alum-formulated vaccine. Studies showed that TNF-α is important in controlling FMD infection. A study by Su et al. showed that the injection of FMD vaccine and TNF-α causes increased immune response (29). As an adjuvant, previous studies on the alum/NLX mixture showed higher Th1 and sometimes Th2 immune responses in combination with various vaccines (26-28, 30). Here, we could not achieve sharp immune responses in the mice immunized with the mixture adjuvant as compared with those immunized with the routine vaccine; the nature of the antigen may be involved in such results.

In the next, real-time PCR was performed to determine the expression profile of IL-12, IL-17 and IFN-γ cytokines. Our findings showed a significant increase in the expression level of IL-12, IL-17 and, IFN-γ mRNAs in mice immunized with the vaccine formulated with the alum/NLX mixture as compared with alum-based vaccine. IL-12 is an inflammatory cytokine that induces the Th1 response (8). A significant increase in the IL-12 secretion by administration of FMD-alum/NLX vaccine candidate was able to induce Th1 responses and can help thorough clearance of FMD infection. In addition, mice immunized with the vaccine formulated with the alum/NLX mixture adjuvant led to increased IL-17 mRNA expression compared to those immunized with the vaccine developed with alum. IL-17 is the signature cytokine produced by Th17 cells. Th17 cells play a prominent role in the infections and lead to recruitment of neutrophils and macrophages into the site of infection (28, 31, 32). In addition, FMD-alum/NLX vaccine formulation led to a significant increase in the IFN-γ mRNA expression. This increased expression in IFN-γ helps eliminate virus-infected cells and prevents the formation of carrier livestock (33). Increased expression of IFN-γ was also observed in a study by Mingala et al. in cattle infected with an inactivated FMD vaccine (34). These results indicated that the alum/NLX mixture could shift the immune responses toward Th1 responses (1, 2, 8, 17, 33, 34).

Humoral immunoassay demonstrated that FMD-alum/NLX and FMD-alum vaccines caused a significant increase in the production of specific total IgG antibodies compared with the other experimental groups. In addition, analysis of antibody isotypes showed that NLX/alum mixture adjuvant-induced the highest level of IgG1 and IgG2a antibodies as compared with other groups. Furthermore, mice immunized with the vaccine formulated in NLX/alum showed increased secretion of IgG2a in comparison with those immunized with the routine vaccine at a borderline. Overall, it seems that the addition of NLX to the vaccine formulation improved humoral immune, as reported in various antigen models (24, 35-37).

In conclusion, results from this study indicated that the alum/NLX mixture could modulate immune responses compared with the routine vaccine. However, other aspects of immune responses, especially the efficacy of this formulation in an experimental viral challenge study, remained to be elucidated.

## Acknowledgment

The authors would like to thank Dr. Morteza Taghizadeh, Dr. Masoud Moghadampour, and Mr. Mohammad Choopani for technical assistance.

## References

.1 Sacerdote P, Gaspani L, Panerai AE. The opioid antagonist naloxone induces a shift from type 2 to type 1 cytokine pattern in normal and skin□grafted mice. Annals of the New York Academy of Sciences. 2000;917(1):755–63.

.2 San Secondo D. Endogenous opioids modulate allograft rejection time in mice: possible relation with Th1/Th2 cytokines. Clinical & Experimental Immunology. 1998;113(3):465–9.

.3 Wernery U, Kinne J. Foot and mouth disease and similar virus infections in camelids: a review. Rev Sci Tech..18–907:(3)31;2012

.4 Kitching R. Global epidemiology and prospects for control of foot-and-mouth disease. Foot-and-Mouth Disease Virus: Springer; 2005. p. 133–48.

.5 Li KB, Liang JP, editors. Assess the Effect of the Foot and mouth disease type O inactivated vaccine (O/MYA98/BY/2010 strain) for pigs. Advanced Materials Research; 2012: Trans Tech Publ.

.6 Zibaei S. purification and preparation of specific antibody against 146 S antigen of FMD virus current serotypes in Iran. 2010.

.7 Brooksby J. Portraits of viruses: foot-and-mouth disease virus. Intervirology. 1982;18(1-2):1–23.

.8 Wang W, Singh M. Selection of adjuvants for enhanced vaccine potency. World Journal of Vaccines. 2011;1(02):33.

.9 Cartwright B, Chapman W, Sharpe R. Stimulation by heterotypic antigens of foot-and-mouth disease virus antibodies in vaccinated cattle. Research in veterinary science. 1982;32(3):338–42.

.10 Coffman RL, Sher A, Seder RA. Vaccine adjuvants: putting innate immunity to work. Immunity. 2010;33(4):492–503.

.11 Seubert A, Monaci E, Pizza M, O’Hagan DT, Wack A. The adjuvants aluminium hydroxide and MF59 induce monocyte and granulocyte chemoattractants and enhance monocyte differentiation toward dendritic cells. The journal of Immunology. 2008;180(8):5402–12.

.12 Cao Y. Adjuvants for foot-and-mouth disease virus vaccines: recent progress. Expert review of vaccines. 2014;13(11):1377–85.

.13 Barnett P, Keel P, Reid S, Armstrong R, Statham R, Voyce C, et al. Evidence that high potency foot-and-mouth disease vaccine inhibits local virus replication and prevents the ‘carrier’state in sheep. Vaccine. 2004;22(9-10):1221–32.

.14 Xu Y, Fu L, Jiang C, Qin Y, Ni Y, Fan J. Naloxone inhibition of lipopolysaccharide-induced activation of retinal microglia is partly mediated via the p3 8mitogen activated protein kinase signalling pathway. Journal of International Medical Research. 2012;40(4):1438–48.

.15 Lee K, Won HY, Bae MA, Hong J-H, Hwang ES. Spontaneous and aging-dependent development of arthritis in NADPH oxidase 2 deficiency through altered differentiation of CD11b+ and Th/Treg cells. Proceedings of the National Academy of Sciences. 2011;108(23):9548–53.

.16 Shebl FM, Yu K, Landgren O, Goedert JJ, Rabkin CS. Increased levels of circulating cytokines with HIV-related immunosuppression. AIDS research and human retroviruses. 2012;28(8):809–15.

.17 Wijesundara DK, Jackson RJ, Tscharke DC, Ranasinghe C. IL-4 and IL-13 mediated down-regulation of CD8 expression levels can dampen anti-viral CD8+ T cell avidity following HIV-1 recombinant pox viral vaccination. Vaccine. 2013;31(41):4548–55.

.18 Velashjerdi Farahani S, Reza Aghasadeghi M, Memarnejadian A, Faezi S, Shahosseini Z, Mahdavi M. Naloxone/alum mixture a potent adjuvant for HIV-1 vaccine: induction of cellular and poly-isotypic humoral immune responses. Pathogens and global health. 2016;110(2):39–47.

.19 Schmittgen TD, Livak KJ. Analyzing real-time PCR data by the comparative C T method. Nature protocols. 2008;3(6):1101.

.20 Levast B, Awate S, Babiuk L, Mutwiri G, Gerdts V. Vaccine potentiation by combination adjuvants. Vaccines. 2014;2(2):297–322.

.21 Gupta RK, Rost BE, Relyveld E, Siber GR. Adjuvant properties of aluminum and calcium compounds. Vaccine design: Springer; 1995. p. 229–48.

.22 Brewer JM, Conacher M, Satoskar A, Bluethmann H, Alexander J. In interleukin□4□deficient mice, alum not only generates T helper 1 responses equivalent to Freund’s complete adjuvant, but continues to induce T helper 2 cytokine production. European journal of immunology. 1996;26(9):2062–6.

.23 Chung W, Liao P, Chen S, Yang P, Lin Y, Jong M, et al. Optimization of foot-and-mouth disease vaccination protocols by surveillance of neutralization antibodies. Vaccine. 2002;20(21-22):2665–70.

.24 Yasaghi M, Mahdavi M. Potentiation of human papilloma vaccine candidate using naloxone/alum mixture as an adjuvant: increasing immunogenicity of HPV-16E7d vaccine. Iranian journal of basic medical sciences. 2016;19(9):1003.

.25 Amini Y, Tebianian M, Mosavari N, Fasihi Ramandi M, Ebrahimi SM, Najminejad H, et al. Development of an effective delivery system for intranasal immunization against Mycobacterium tuberculosis ESAT-6 antigen. Artif Cells Nanomed Biotechnol. 2017;45(2):291–6.

.26 Jazani NH, Parsania S, Sohrabpour M, Mazloomi E, Karimzad M, Shahabi S. Naloxone and alum synergistically augment adjuvant activities of each other in a mouse vaccine model of Salmonella typhimurium infection. Immunobiology. 2011;216(6):744–51.

.27 Deepak S, Kottapalli K, Rakwal R, Oros G, Rangappa K, Iwahashi H, et al. Real-time PCR: revolutionizing detection and expression analysis of genes. Current genomics. 2007;8(4):234–51.

.28 Tesmer LA, Lundy SK, Sarkar S, Fox DA. Th17 cells in human disease. Immunological reviews. 2008;223(1):87–113.

.29 Su B, Wang J, Wang X, Jin H, Zhao G, Ding Z, et al. The effects of IL-6 and TNF-α as molecular adjuvants on immune responses to FMDV and maturation of dendritic cells by DNA vaccination. Vaccine. 2008;26(40):5111–22.

.30 Harandi AM, Medaglini D, Shattock RJ, convened by EUROPRISE WG. Vaccine adjuvants: a priority for vaccine research. Vaccine. 2010;28(12):2363–6.

.31 Curtis MM, Way SS. Interleukin-17 in host defence against bacterial, mycobacterial and fungal pathogens. Immunology. 2009;126(2):177–85.

.32 Najminejad H, Kalantar SM, Mokarram AR, Dabaghian M, Abdollahpour-Alitappeh M, Ebrahimi SM, et al. Bordetella pertussis antigens encapsulated into N-trimethyl chitosan nanoparticulate systems as a novel intranasal pertussis vaccine. Artif Cells Nanomed Biotechnol. 2019;47(1):2605–11.

.33 Rizk SA, EL-Din WMG, Mahdy SE, Ibrahim EE-S, Moha H. Gamma Interferon Assay for Cellular Immune Response in Cattle Vaccinated with FMD Vaccine Adjuvanted with Different Montanide Oils. Global Journal of Medical Research. 2015.

.34 Mingala CN, Konnai S, Venturina FA, Onuma M, Ohashi K. Quantification of water buffalo (Bubalus bubalis) cytokine expression in response to inactivated foot-and-mouth disease (FMD) vaccine. Research in veterinary science. 2009;87(2):213–7.

.35 Jamali A, Mahdavi M, Hassan ZM, Sabahi F, Farsani MJ, Bamdad T, et al. A novel adjuvant, the general opioid antagonist naloxone, elicits a robust cellular immune response for a DNA vaccine. International immunology. 2009;21(3):217–25.

.36 Jamali A, Mahdavi M, Shahabi S, Hassan ZM, Sabahi F, Javan M, et al. Naloxone, an opioid receptor antagonist, enhances induction of protective immunity against HSV-1 infection in BALB/c mice. Microbial pathogenesis. 2007;43(5-6):217–23.

.37 Ahmadzadeh A, Mahdavi SM, Karimi P, Nikzamir A. Naloxone inhibits human serum albumin Glycation. Journal of Paramedical Sciences. 2014;5(3).

